# The fecal microbiota transplantation from drug-naïve schizophrenia patients distinctively changes gut microbiome and metabolic profiles in male and female mice

**DOI:** 10.1101/2025.09.13.675957

**Authors:** Raghunath Singh, Sandra Pereira, Kristoffer Panganiban, Laurie Hamel, David C. Wright, Thomas D. Prevot, Daniel Mueller, Gary Remington, Sri Mahavir Agarwal, Premysl Bercik, Elena F Verdu, Giada De Palma, Margaret K Hahn

**Author notes:** Shared Senior Authors (Giada De Palma, Margaret K Hahn). Corresponding Author: Margaret K Hahn, MD, PhD, FRCPC Clinician-Scientist Lead, Mental Health and Metabolism Clinic Complex Care and Recovery, Schizophrenia Division Center for Addiction and Mental Health (CAMH) Professor, University of Toronto, Department of Psychiatry Kelly and Michael Meighen Chair in Psychosis Prevention.

## Abstract

**Background:** Emerging evidence suggests a role for the gut microbiome in schizophrenia (SCZ) and antipsychotic-induced metabolic perturbations. Using human fecal microbiota transplantation (FMT) in mice, this study investigated the role of gut microbiome in metabolic changes related to SCZ and antipsychotic (olanzapine) treatment.

**Methods:** 5-6 weeks old germ-free NIH Swiss mice of both sexes received microbiota from either SCZ patients (SCZ-FMT) or healthy controls (HC-FMT) followed by a diet with or without olanzapine for six-weeks. Food intake and body weight were monitored weekly, and an intraperitoneal glucose tolerance test and open field test were performed. Serum glucose, and insulin were measured. Gut microbiome characterization and short-chain fatty acids (SCFAs) quantification were performed in the cecal samples using 16S rRNA gene sequencing and gas chromatography-mass spectrometry, respectively.

**Results:** Olanzapine treatment decreased the locomotor activity in the open field test, irrespective of sex or microbiota. Female SCZ-FMT recipient mice exhibited insulin resistance compared to HC-FMT, irrespective of olanzapine treatment. Female SCZ-FMT mice showed significantly lower alpha-diversity compared to HC-FMT, whereas olanzapine treatment increased alpha-diversity. SCZ-FMT and olanzapine treatment differentially altered the microbial abundances, and metabolic pathways in male and female mice. Interestingly, cecal SCFAs, mainly acetate levels, were significantly decreased in female SCZ-FMT mice compared to HC-FMT, while olanzapine treatment increased acetate levels in male mice. Both male and female SCZ-FMT mice showed elevated levels of isovaleric acid compared to HC-FMT.

**Conclusion:** These preliminary findings suggest that gut microbiome could be a predisposing factor contributing to the intrinsic risk of developing type 2 diabetes associated with SCZ in females.

Graphical abstract

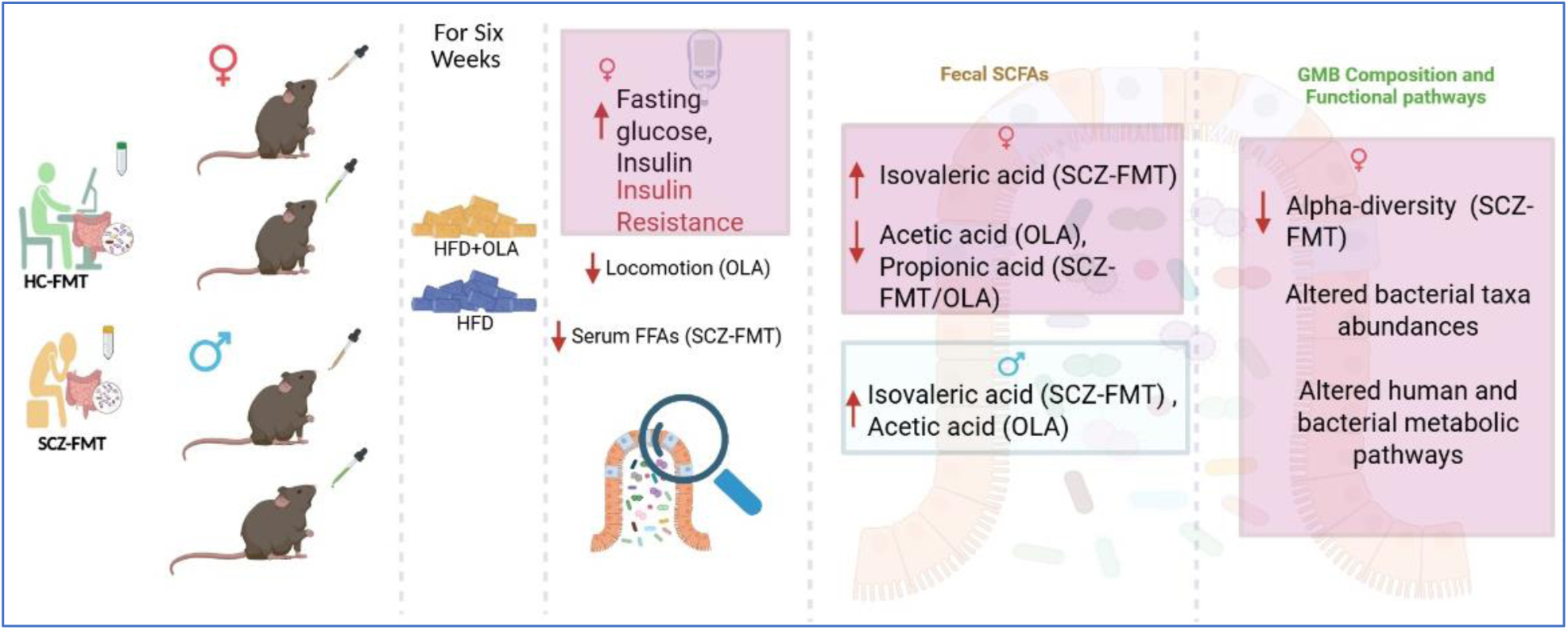

## Introduction

Schizophrenia (SCZ) is a severe and complex mental illness affecting nearly 1% of the world population and is among the top 10 global causes of disability [1]. Individuals with SCZ have a reduced lifespan, experiencing a mortality rate double that of the overall population [2], largely attributable to cardiometabolic disorders such as obesity, and type 2 diabetes (T2D) [3]. SCZ patients have a 33% higher risk of T2D compared to the general population [4]. Before the discovery of chlorpromazine in the 1950s, diabetes was historically linked to mental disorders. A metanalysis of 1137 participants links first-episode psychosis, insulin resistance, and impaired glucose tolerance [5]. Childhood fasting insulin levels and body mass index were linked to psychosis and depression in young adults over time [6], suggesting the intrinsic risk of T2D in patients with SCZ. Antipsychotic (AP) medications are very successful in reducing symptoms of psychosis. Yet, they are commonly linked to various cardiometabolic side effects such as significant weight gain, T2D development and progression, and liver disease [7]. Along with others [8–10], we have also shown that APs acutely disrupt glucose metabolism by dysregulating insulin’s action on the brain [11, 12] to suppress endogenous glucose production, and disrupt hypothalamic nutrient sensing in vivo [13, 14], and in vitro [15, 16]. Nevertheless, these findings failed to consider the intrinsic risk of T2D development among individuals with SCZ.

The gut microbiota (GMB) has recently been recognized as potentially playing a role in SCZ [17].. Altered GMB composition has been associated with psychiatric disorders such as major depression, bipolar disorder, SCZ, and anxiety [18]. In particular, altered GMB has been linked to SCZ pathogenesis [18–20], symptoms severity, treatment response [20, 21], and treatment resistance [22]. Patients with ultra-high risk of SCZ have shown significantly different GMB composition compared to patients with high risk of SCZ and healthy controls, and this was associated with elevated short-chain fatty acids (SCFAs) biosynthesis pathways [23]. SCFAs, GMB-derived metabolites, such as isovaleric acid are associated with SCZ symptoms such as negative symptoms, cognition impairments, and subclinical inflammation [24, 25].

Alongside playing a role in the pathogenesis of SCZ, the GMB has been implicated in AP-associated metabolic alterations [17] with evidence arising from clinical [26–31], and rodent studies [32–35]. Rodent studies have also shown that fecal microbiota transplantation (FMT) from SCZ patients (SCZ-FMT) into germ-free or antibiotics-treated mice induces SCZ-type behavioral and molecular phenotypes [20, 36–38]. To date, however, no causal link has been established between SCZ and the intrinsic risk of developing T2D. Therefore, this study aimed at investigating the causal role of GMB in the relationship between T2D and SCZ, using SCZ-FMT in male and female germ-free mice.

## Methods

### Fecal microbiota transplantation (FMT)

Fecal donors were selected from our ongoing clinical trial protocol (clinicaltrials.gov: NCT03414151). FMTs were prepared from AP-naïve SCZ patients (SCZ-FMT) who gained >5% body weight after 12 weeks of antipsychotic treatment (Patient 1 12.3%, patient 2-9.4%), and age-sex-BMI matched healthy controls (HC-FMT).

### Animals, diets and treatments

Germ-free NIH Swiss mice (5-6 weeks old) were provided by the Axenic Gnotobiotic Unit (AGU) of McMaster University. Germ-free mice (n = 87, 39 females) were colonized with diluted human fecal samples (1:10 in saline) and housed for 4 weeks in sterilized ventilated racks on a 12-hour light/12-hour dark cycle with free access to food and water, as described previously [39, 40]. At the age of 10-12 weeks, mice were given an irradiated high-fat diet (HFD) as a control diet (45% kcal, Con) or HFD + Olanzapine (OLA) for six additional weeks in a level-2 containment facility in individually ventilated cages. OLA (50 mg/kg of the diet) was compounded in HFD to achieve an accelerated model of AP-induced metabolic and GMB alterations in mice [32, 41, 42]. Weekly food intake and body weight were monitored. Weight gain was expressed as percentage weight change calculated from initial and final body weight using the formula: [% weight change = ((final-initial)/initial)*100]. During the last week, animals were subjected to an intraperitoneal glucose tolerance test (IPGTT), followed by the open field test to assess locomotor activity [43]. For the IPGTT, overnight food-restricted mice were given glucose (2g/kg) intraperitoneally, and glucose was measured using a glucometer (AccuCheck). Circulating concentrations of insulin measured using ELISA kits (Alpco). Homeostatic model assessment for insulin resistance (HOMA-IR), and quantitative insulin sensitivity check index (QUICKI) were calculated for the assessment of insulin sensitivity.

### Cecal SCFA analysis

Cecal samples were acidified by adding a weight equivalent of 3.7% HCl. The acidified samples were extracted three times with propyl formate containing butyric acid-d_7_ internal standard, and a 60 µL extract aliquot was derivatized with 25 µL N-tert-butyldimethylsilyl-N-methyltrifluoroacetamide (MTBSTFA) at 40°C for 1 hour then analyzed by chromatography mass spectrometry (GC-MS). Acetic acid, propionic acid, isobutyric acid, butyric acid, isovaleric acid, and pentanoic acid were quantified and reported as nmol/mg of sample. The calibration curves for all analytes were linear over the range of interest, with R^2^ > 0.99. A combined pooled sample was employed for reproducibility quality control checks and separated into four aliquots (2 unspiked and 2 spiked for recovery testing) [40].

### 16S rRNA gene sequencing

Fecal microbiota analysis was done using 16S rRNA gene sequencing according to previous described methods [39, 40]. Briefly, mice fecal and cecal samples were collected at the end of the experiment and immediately frozen at –80°C. Total genomic DNA was extracted from the stool samples as previously described [44]. Amplification of the V3-V4 region of the 16S rRNA gene, and Illumina sequencing were performed as previously described [44, 45]. Data were analyzed following the pipelines of Divisive Amplicon De Noising Algorithm 2 (dada2) [46] and Phyloseq packages (1.28) [47] for R (3.6.1). Taxonomic assignments were performed using the RDP classifier [48] with the Silva small subunit Ref. NR99 138.1 database [49, 50] (2020) training set. Samples with <1000 reads were removed from the analysis, resulting in a median frequency of 58,238 reads per sample. Analyses were done using plugins available in Quantitative Insights into Microbial Ecology 2 QIIME2 [51] and Maaslin2 [52]. Metagenomic functional content was predicted from 16S rRNA gene profile of samples using PICRUSt 2.0 (v. 2.4.1) [53], analyzed using Maaslin2 [52] and visualized using Stamp [54]. All results were corrected for multiple comparisons, allowing 5% false discovery rate.

## Statistical analysis

Statistical analysis was carried out using GraphPad Prism 10.3.0 (GraphPad Software, La Jolla, CA, USA), and R (version 4.1.1). Statistical comparisons were performed using Two-way ANOVA followed by Uncorrected Fisher’s LSD post-test. *Post-hoc* analysis was performed when there was a statistical interaction between disease (SCZ) and treatment (OLA) (*P* < 0.05). Repeated measures ANOVA was used to analyze parameters at multiple time points in IPGTT, across groups. *P* <0.05 was considered statistically significant.

## Data availability statement

All supporting data are available within the article and its Supporting Data Values file. 16S rRNA gene sequencing data generated in this study will be deposited in the National Center for Biotechnology Information (NCBI), Sequence Read Archive (SRA).

## Results

### The SCZ microbiota and OLA treatment do not affect body weight and food intake

Weight gain and average daily food intake were not affected by the microbiota (SCZ or HC) or by olanzapine (OLA) treatment in both humanized male and female mice (Fig. 1B, G, E, and F). However, OLA treatment significantly decreased locomotion (distance traveled in the open field test), regardless of sex or microbiota (*P* < 0.05 in female, and *P* < 0.01 in male) (Fig. 1D, and G).

**Figure 1.**
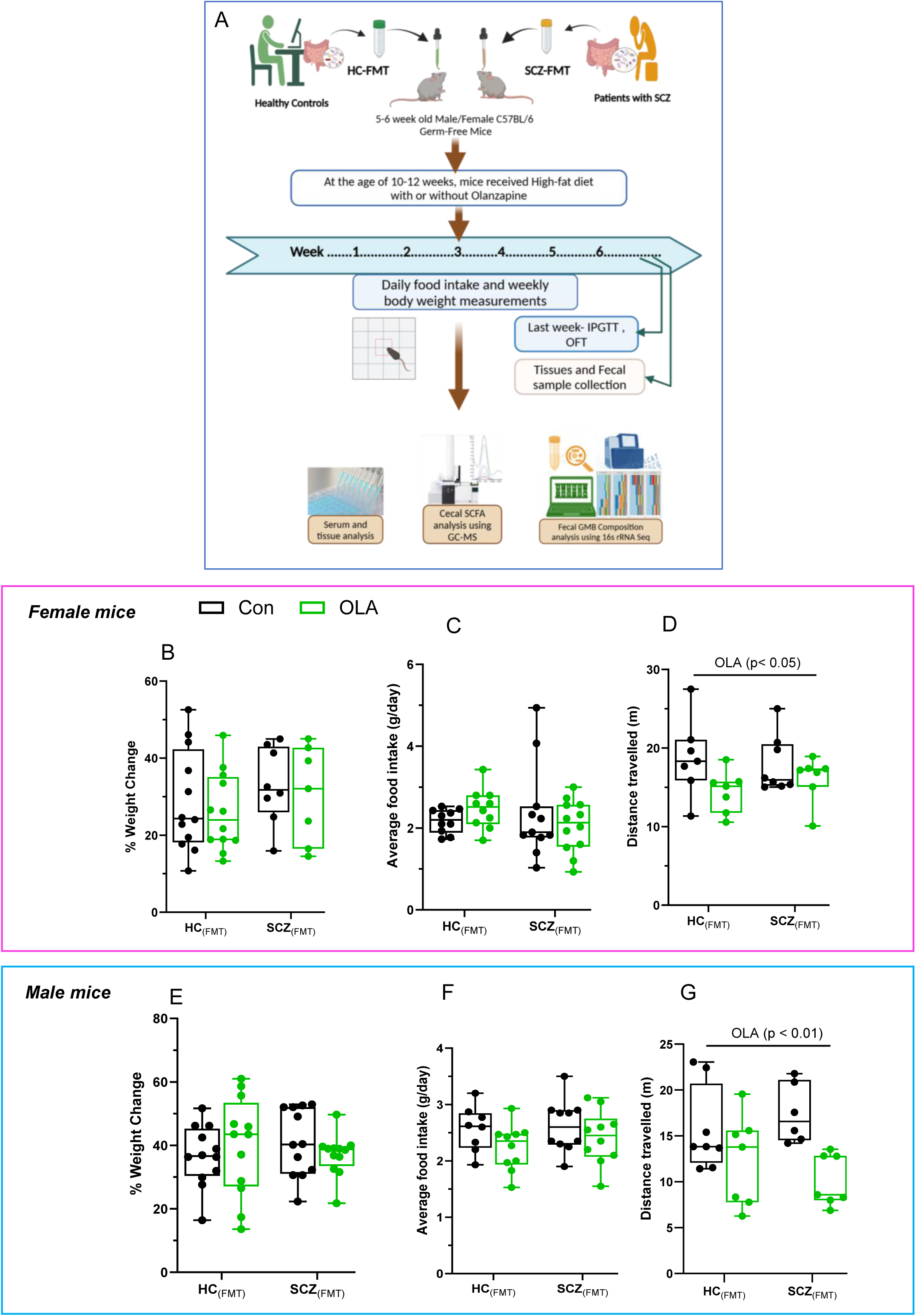
Effect SCZ microbiota and OLA treatment on body weight, food intake and locomotion. A) Schematic representation of the experimental design. B-D) Effect of SCZ_(FMT)_ and OLA treatment on % body weight change (calculated from initial and final body weights), average food intake (g/day/mouse), and distance traveled (locomotion) in open filed test (OFT) in female mice (n= 7-12). E-G) Effect of SCZ_(FMT)_ and OLA treatment on % body weight change, average food intake, and distance traveled in OFT in male mice (n= 8-12). Data were analyzed with two-way ANOVA followed by Uncorrected Fisher’s LSD test for multiple comparisons when a statistically significant interaction was observed. Data are presented as interleaved box and whisker plots with whiskers min-max. *P* < 0.05 was considered statistically significant. P values marked with “OLA” refer to the effect of OLA treatment vs Control diet. Abbreviations: HC_(FMT)_ and SCZ_(FMT)_-germ-free mice receiving fecal transplant from HC and SCZ patients, respectively. Con-control diet, OLA-Olanzapine diet.

### The SCZ microbiota induces insulin resistance only in female mice

Six weeks of OLA treatment significantly increased fasting blood glucose in female mice, irrespective of the microbiota (HC or SCZ) (*P* < 0.05) (Fig. 2A), while no significant difference was observed in male mice (Fig. 2H). Gut microbiota from SCZ patients significantly showed deleterious effect on insulin sensitivity, particularly in female mice. SCZ-FMT significantly increased fasting serum insulin levels compared to HC-FMT in female mice (*P* < 0.01), resulting in higher HOMA-IR (*P* < 0.01) and lower QUICK index (*P* < 0.01), which are the indices to measure insulin sensitivity (Fig. 2B-D). Male mice did not show any significant difference (Fig. I-K). Interestingly, female SCZ-FMT recipient mice showed significantly decreased area under the curve during the IPGTT compared to HC-FMT (P <0.05). This effect was irrespective of OLA treatment (Fig. 2E-F). In male mice, insulin sensitivity and glucose tolerance remain intact (Fig. 2G-L).

**Figure 2.**
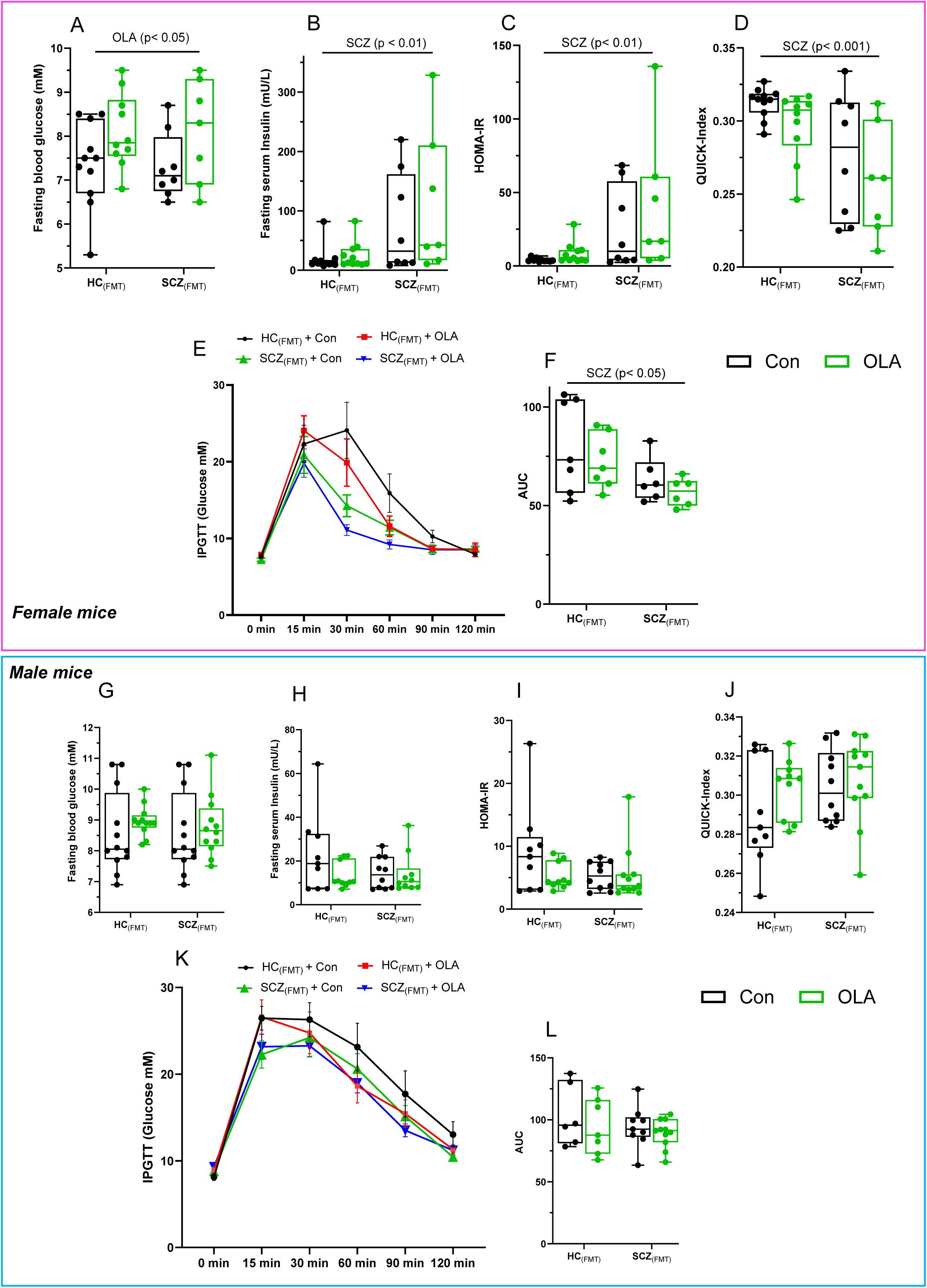
Effect of SCZ microbiota and OLA treatment on metabolic parameters. A-D) Effect of SCZ_(FMT)_ and OLA treatment on fasting blood glucose and insulin levels, HOMA-IR, and QUICK-Index in female mice (n= 7-11). E-F) Effect of SCZ_(FMT)_ and OLA treatment intraperitoneal glucose tolerance test (IPGTT) and respective area under the curve (AUC) in female mice. G-L) Effect of SCZ_(FMT)_ and OLA treatment on fasting blood glucose and insulin levels, HOMA-IR, and QUICK-Index in male mice (n= 9-10). L-N) Effect of SCZ_(FMT)_ and OLA treatment on IPGTT, and respective AUC in male mice (n= 8-10). SCZ is showing effect of SCZ_(FMT)_ (vs HC_(FMT)_), and OLA is showing effect of OLA treatment (vs Con). Data were analyzed with two-way ANOVA followed by Uncorrected Fisher’s LSD test for multiple comparisons when a statistically significant interaction was observed. Repeated measures two-way ANOVA was used to analyze glucose concentration at different time points in IPGTT. Data are presented as interleaved box and whisker plots with whiskers min-max. *P* < 0.05 was considered statistically significant. P values marked with “OLA” refer to the effect of OLA treatment vs Con; P values marked with “SCZ” refer to the effect of SCZ_(FMT)_ vs HC_(FMT)_. Abbreviations: HC_(FMT)_ and SCZ_(FMT)_-germ-free mice receiving fecal transplant from HC and SCZ patients, respectively. Con-Control diet, OLA-Olanzapine diet; HOMA-IR-homeostatic model assessment for insulin resistance; QUICKI-Quantitative Insulin sensitivity Check Index.

### The SCZ microbiota and OLA treatment alter cecal short-chain fatty acid level

To determine the functional impact of the gut microbiota from SCZ patients in humanized mice, we measured cecal SCFAs levels using GC-MS. In male mice, OLA treatment significantly increased cecal acetic acid levels, irrespective of the microbiota (HC or SCZ) (*P* < 0.01) (Fig. 3F), while no significant difference was observed in the female mice (*P* = 0.0723). Interestingly, OLA treated female SCZ-FMT mice showed decreased acetic acid levels compared to OLA treated HC-FMT mice (*P* < 0.05, Fig. 3A). The gut microbiota from SCZ patients increased cecal isovaleric acid levels significantly compared to HC microbiota in both female and male mice (*P* < 0.05, and *P* < 0.01, respectively; Fig. 3D and I). Sex-independent elevated levels of isovaleric acid in SCZ-FMT mice could suggest disease (SCZ) specific alterations. Finally, OLA treatment significantly decreased cecal propionic acid levels in SCZ-FMT female mice only (*P* < 0.05) (Fig. 3E, and J).

**Figure 3.**
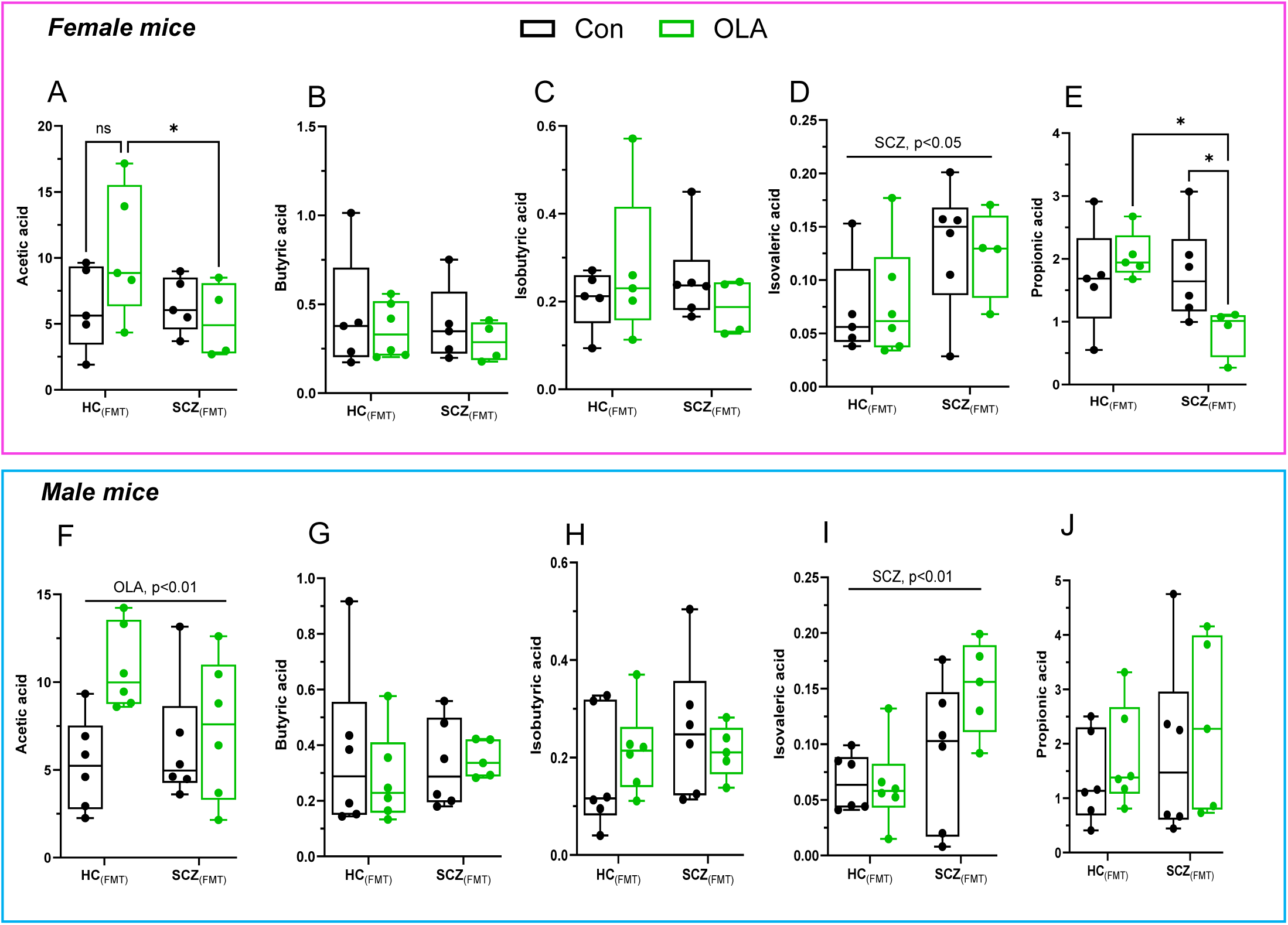
Effect SCZ microbiota and OLA treatment on cecal SCFAs levels. A-E) Cecal levels of SCFAs in female mice (acetic, butyric, isobutyric, isovaleric, and propionic acid). SCFAs are measured using gas chromatography-mass spectroscopy (GC-MS) and expressed as nmol/mg of sample. F-J) Cecal levels of SCFAs (nmol/mg of sample) in male mice. SCZ shows effect of SCZ_(FMT)_ (vs HC_(FMT)_), Data were analyzed with two-way ANOVA followed by Uncorrected Fisher’s LSD test for multiple comparisons when a statistically significant interaction was observed. Data are presented as interleaved box and whisker plots with whiskers min-max. n = 4–6 mice/group. *P* < 0.05 was considered statistically significant. P values marked with “OLA” refer to the effect of OLA treatment vs Con; P values marked with “SCZ” refer to the effect of HC_(FMT)_ vs SCZ_(FMT)_. Abbreviations: HC_(FMT)_ and SCZ_(FMT)_-germ-free mice receiving fecal transplant from HC and SCZ patients, respectively. Con-control diet, OLA-Olanzapine diet. * = *P* < 0.05 (HC_(FMT)_+OLA vs SCZ_(FMT)_+OLA; or SCZ_(FMT)_ vs SCZ_(FMT)_+OLA).

### The gut microbiota composition differs between SCZ-FMT and HC-FMT recipient mice in a sex-dependent manner

To validate the fidelity of FMT from human donors to recipient mice, we plotted the beta-diversity of both donors and recipient mice (Fig.4A). Despite some minor differences in microbiota composition between mouse recipients and human donors (supplementary figure1), similar to the previous reports [39, 40], SCZ and HC microbiota profiles were transferred into gnotobiotic mice. The microbial profiles clustered separately by microbiota (Permanova, *P* = 0.001) and closer to their respective donors. We also observed a significant effect of sex in the clustering (Permanova*, P* = 0.024). However, there was no effect of OLA treatment on the microbiota composition of SCZ-FMT or HC-FMT mice (Fig. 4B-C).

**Figure 4:**
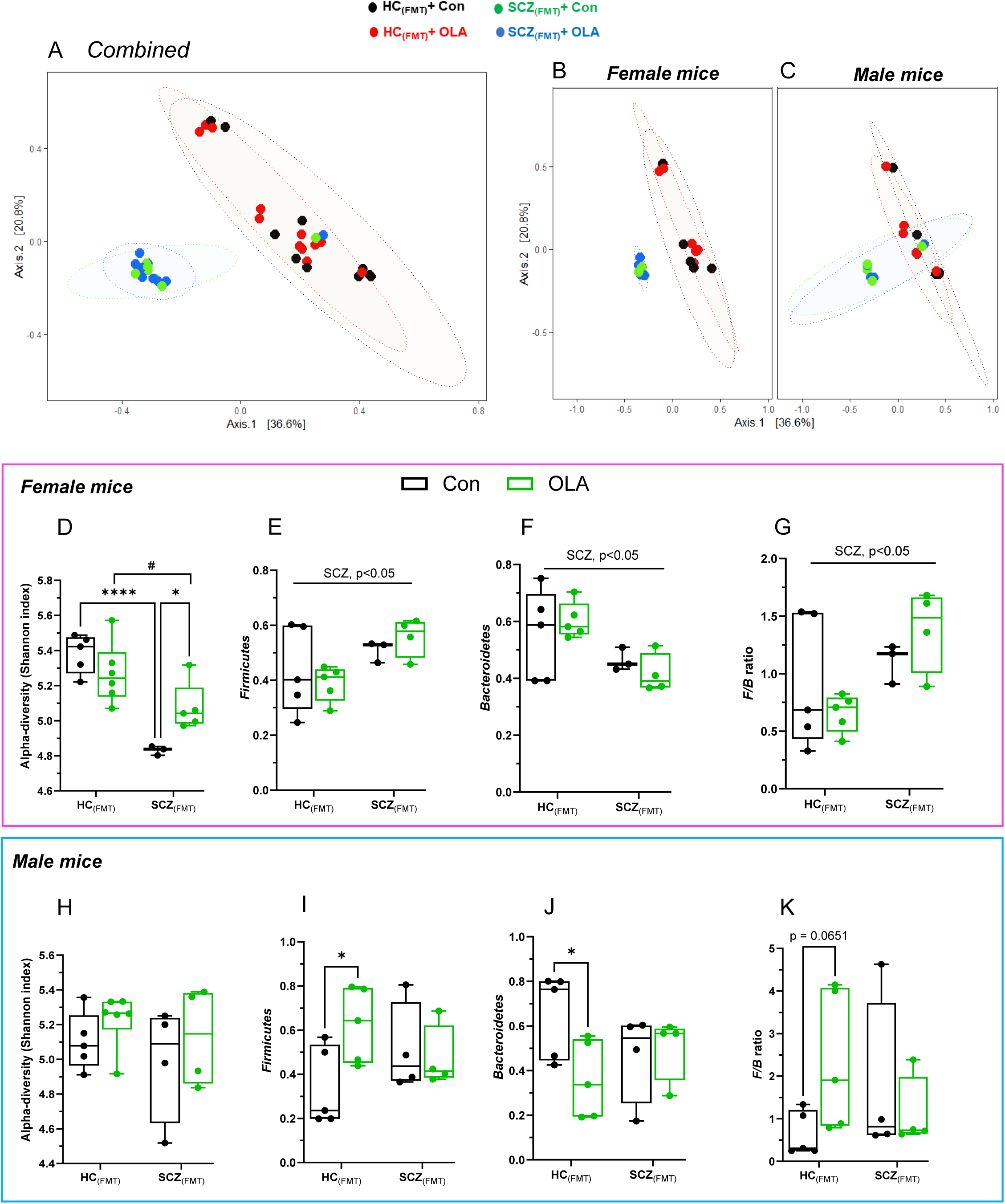
Gut microbiota composition in donor (HC or SCZ) and fecal microbiota recipient mice. A) Principal coordinates analysis (PCoA) ordination plots of the Bray Curtis dissimilarities matrix computed on rarefied data showing ‘beta-diversity’ in FMT recipient mice (combined, male and female). Black/red circles show healthy control (HC_(FMT)_) recipients mice treated with Con, and OLA, respectively. Green/blue circles show schizophrenia (SCZ_(FMT)_) recipients treated with Con, and OLA, respectively. Community composition was compared by ADONIS2 with 999 permutations. Ellipses indicate 95% confidence intervals and were generated as a distribution around a centroid for each group based on ADONIS2. B-C) 2D PCoA plots of Bray Curtis dissimilarities showing ‘beta-diversity’ in different treatment groups in female and male mice. D, H) Microbial (alpha)-diversity (Shannon diversity index) in female and male mice. E-G) relative abundances of Firmicutes, Bacteroidetes and their ratio (F/B ratio) in female mice. I-K) relative abundances of Firmicutes, Bacteroidetes and F/B ratio in male mice. Microbial diversity data were analyzed with two-way ANOVA followed by Uncorrected Fisher’s LSD test for multiple comparisons. Data are presented as interleaved box and whisker plots with whiskers min-max. n = 3-5 mice/group. *P* < 0.05 was considered statistically significant. \*\*\*\**P* < 0.0001 (HC_(FMT)_ vs SCZ_(FMT)_, ^#^*P* < 0.05 (HC_(FMT)_+OLA vs SCZ_(FMT)_+OLA), and \**P* < 0.05 (SCZ_(FMT)_/HC_(FMT)_ + Con vs SCZ_(FMT)_/HC_(FMT)_ +OLA). Abbreviations: HC_(FMT)_ and SCZ_(FMT)_-germ-free mice receiving fecal transplant from HC and SCZ patients, respectively. Con-control diet, OLA-Olanzapine diet, and F/B ratio-Firmicutes/Bacteroidetes ratio.

When we analyzed the microbial diversity of SCZ-FMT and HC-FMT mice, the gut microbiota of SCZ patients induced lower alpha-diversity (Shannon diversity index) in SCZ-FMT recipient female mice in comparison to HC-FMT (*P* < 0.0001) (Fig. 4D). OLA treatment, however, significantly increased microbial diversity in SCZ-FMT female mice (*P* < 0.05), but not to the levels of HC-FMT mice. No significant differences were observed in male mice (Fig. 4H). We then analyzed the relative abundance of two taxa previously linked to metabolism alterations, such as Firmicutes, Bacteroidetes and their ratio (F/B ratio). SCZ-FMT female mice, and not male mice, presented with an increase in Firmicutes, a decrease in Bacteroidetes relative abundance and an increased F/B ratio (Fig. 4E-G). In HC-FMT male mice, however, OLA treatment significantly increased the F/B ratio (*P* = 0.0651) compared to Con (Fig. 4I-K).

At taxonomical level, the gut microbiota composition of SCZ-FMT and HC-FMT mice was significantly different and influenced by OLA treatment in a sex-dependent manner (Fig. 5A-B). Several bacterial genera (identified by ANCOM-BC analysis) were found to be altered in both female and male SCZ-FMT mice in comparison to HC-FMT (Fig. 5C-D). Interestingly, we found that the gut microbiota composition was primarily influenced by the source of microbiota (SCZ vs. HC) rather than OLA treatment.

**Figure 5.**
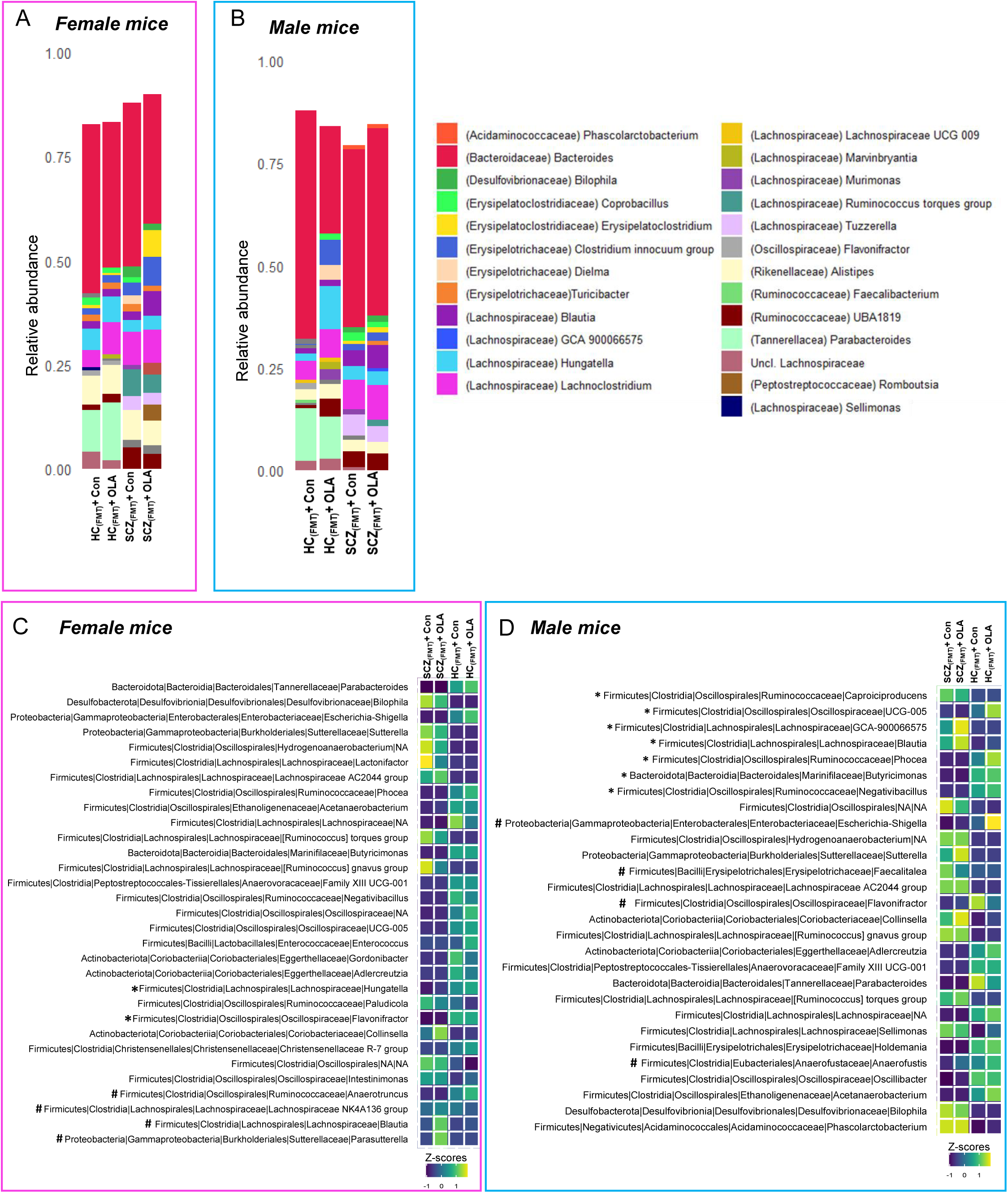
Differential abundance of bacterial taxa in male and female mice receiving SCZ-FMT and OLA treatment. A and B) Taxonomic bar plots at the family level Taxonomical bar plots showing relative abundance at family level in female and male mice. n = 3–5 mice/group. C and D) Heatmaps of all the differentially abundant taxa (aggregated at genus level) in mice evaluated using ANCOM-BC. Statistical significance on differentially abundant taxa between groups was corrected for multiple comparisons using FDR (q < 0.05). Color intensity on heatmaps was generated by using z-scores calculated based on relative abundance of each bacterial taxa. All bacterial genera shown in the heatmap were differentially present in both the SCZ_(FMT)_ mice groups (Con or OLA), except for those marked with **\*** that were significantly different only in the SCZ_(FMT)_ + Con group (vs the reference group HC_(FMT)_ + Con), and those marked with **#** that were significantly different only in the SCZ_(FMT)_ + OLA group (vs the reference group HC_(FMT)_ + Con). *P* < 0.05 was considered significant.

In female SCZ-FMT recipients presented with a significant increase in the relative abundance of *Hungatella* spp. and *Flavonifractor* spp. when on a control diet but presented with a significant increase in the relative abundance of *Blautia* spp. and *Parasutterella* spp., and a significant decrease in the relative abundance of *Anaerotruncus* spp. when administered OLA treatment (Fig. 5C). Conversely, SCZ-FMT recipient male mice presented with a significant increase in the relative abundance of *Escherichia-Shigella*, *Faecalitalea*, *Flavonifractor*, and *Anaerofustis* spp. when on control diet. OLA treatment increased significantly *Blautia* spp., *Lachnospiraceae* species GCA-900066575, and *Caproiciproducens* spp. in male SCZ-FMT mice, but reduced significantly the relative abundance of *Butyricimonas* spp. and several Clostridia species, such as UCG-005, *Phocea* and *Negativibacillus* (Fig. 5D).

### Effect of SCZ-FMT and OLA treatment functional metabolic pathways

We then specifically investigated the 16S-inferred functional pathways and enzymes that have been previously associated to SCZ, such as tryptophan metabolism (5HT synthesis and degradation), tyrosine metabolism (dopamine synthesis and degradation), GABA-Glutamate synthesis shuttle, histamine, SCFA synthesis and metabolism among others. In female SCZ-FMT mice we found statistically significant differences in genes related to glutamate utilization, while glutamine synthetase (converts glutamate to glutamine) was increased, glutamate decarboxylase (or glutamic acid decarboxylase (GAD), an enzyme that catalyzes the decarboxylation of glutamate to gamma-aminobutyric acid (GABA)), was significantly decreased in female SCZ-FMT mice. In addition, genes related to butyrate synthesis and gluconeogenesis were significantly enriched in SCZ-FMT female mice, and acetyl CoA C-acetyltransferase (synthesis and catabolism of fatty acids), 3-hydroxybutyryl-CoA dehydrogenase (biosynthesis of 3-hydroxybutyrate), and L-lactate dehydrogenase (LDH, conversion of pyruvate to lactate and back) were enriched in female SCZ-FMT mice treated with OLA. Furthermore, in male mice, SCZ-FMT showed enrichment of pathways such as glycolysis II (from fructose 6-phosphate), and super-pathway of glucose and xylose degradation, however, no treatment effect was observed (Fig. 6A, B).

**Figure 6:**
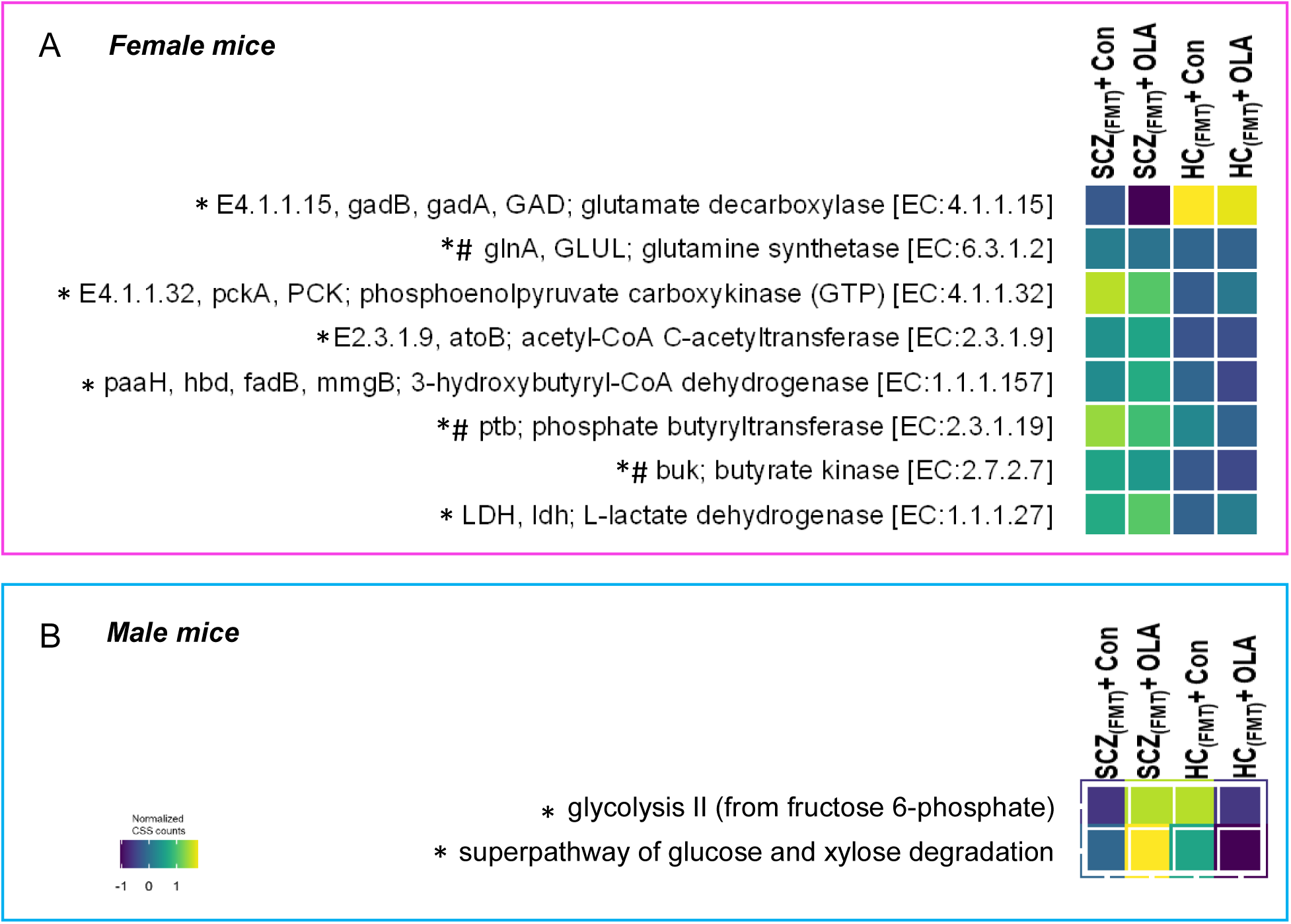
Functional analysis of alteration in potential human or bacterial metabolic pathways. A) Enzymes (EC#) were extracted from the inferred functional profiles of the microbial community based on 16S marker gene data and analyzed across groups. Raw counts were converted into CSS counts to make a heatmap. Maaslin2, and NEGBIN methods were used to analyze count data. B and C) KEGG pathways information was extracted from extracted from the inferred functional profiles of the microbial community based on 16S marker gene data and analyzed across female and male mice. Raw counts were normalized to CSS counts. * denotes a significant difference in the SCZ_(FMT)_ + Con diet group (vs all others), # denotes a significant difference in the SCZ_(FMT)_ + OLA group (vs all others).

## DISCUSSION

Recent preclinical studies have demonstrated the pivotal role of gut microbiota in SCZ pathogenesis. Indeed, fecal microbiota transplantation from SCZ patients (SCZ-FMT) to germ-free or antibiotics-treated mice results in SCZ-like behavioral and molecular phenotypes [20, 36–38]. However, this approach has not been exploited to investigate intrinsic metabolic risks associated with SCZ and AP medication. In the present study, we investigated the role of GMB in metabolic alterations associated with SCZ and OLA (a prototype AP) treatment using human FMT into germ-free mice.

Previous studies found that SCZ-FMT in mice resulted in heightened spontaneous locomotion, quantified by increased distance traveled during open field tests, a potential indicator of schizophrenia-like hyperactivity and positive symptoms [20, 37, 38]. In contrary, we did not find any difference in the locomotor activity in SCZ-FMT groups, but OLA treatment decreased locomotor activity, irrespective of sex or source of microbiota. This is likely due to its sedative property mostly correlated with reduced energy expenditure and consequent weight gain [55–57].

Chronic treatment with antipsychotics, particularly OLA and risperidone, increases food intake and weight gain in conventional female C57BL6 mice [32, 41, 42, 58–60]. OLA treatment is often administered in HFD to achieve an accelerated model of AP-induced metabolic alterations in mice [32, 41, 42]. Nonetheless, it has been shown that the microbiota is necessary to observe the AP-induced weight gain, as female germ-free mice do not gain weight after seven weeks of OLA treatment [32]. Indeed, colonization of germ-free mice and another seven-week treatment with OLA+HFD induced significant weight gain [32]. Here we found that OLA treatment for six weeks did not cause significant weight gain or increased food intake irrespective of sex or microbiota. However, regardless of sex or microbiota, mice gained over 30-45% of their starting weight, which is a significant weight gain resulted from HFD feeding. OLA diet, however, did not induce a greater weight gain than the control diet (HFD) alone in mice receiving SCZ-FMT or HC-FMT.

Even though chronic OLA treatment did not cause hyperphagia and additional weight gain, in comparison to the control HFD, it did induce hyperglycemia in female mice, irrespective of microbiota. This effect was in line with the previous studies showing chronic OLA-induced hyperglycemia in female mice [41, 42, 59]. Surprisingly, male mice were protected against the hyperglycemia caused by OLA, regardless of the microbiota. This effect was also contrary to another study showing only female mice to be protected against acute OLA-induced hyperglycemia, [9, 10]. Our study revealed that only female SCZ-FMT mice after OLA exhibited considerable insulin resistance, characterized by high insulin levels (hyperinsulinemia), increased HOMA-IR, and a lower QUICK-index. To our knowledge, this is first preclinical study suggesting the role of GMB in SCZ-related intrinsic risk of developing insulin resistance using SCZ-FMT in female mice. While in first-episode drug-naïve and chronic SCZ patients, GMB changes and insulin resistance were found to be associated with SCZ symptoms [17, 61, 62] and antipsychotic-induced metabolic alterations [58, 63, 64], we found that only female SCZ-FMT mice had lower glucose levels in the IPGTT. OLA, on the other hand, did not affect insulin resistance and glucose tolerance in IPGTT in male or female mice. Although, these results are in contrast with previous studies [9, 10, 41, 42], it is plausible that other factors (i.e. hyperinsulinemia, sex specific GMB composition, and SCFAs levels to name few) might play a role in the antipsychotic-induced metabolic alterations.

SCFAs, metabolites produced by gut bacteria from complex carbohydrates, are key to communication between the gut and the brain [65, 66]. Most of the SCFAs (such as acetate, propionate, valerate, and butyrate) act through their G-protein coupled receptor which are mainly GPR41, GPR43, and GPR109A, a type of G-protein coupled receptor, also known as free fatty acid receptor 2 (FFAR2), FFAR3, and hydrocarboxylic acid receptor (HCAR2), respectively [65, 67]. SCFAs can enter the systemic circulation and cross the blood-brain barrier (BBB) to regulate pathways related to oxidative stress, metabolism, and neurotransmission, which are all important mechanisms underlying the pathology and symptomatology of SCZ [66, 68]. Propionate has been reported to induce postprandial hyperglycemia in mice and chronic exposure to propionate induces weight gain and insulin resistance in mice and humans [69]. In addition, higher serum levels of acetic acid, acetic acid/propionic acid ratio, has been associated with poorer cognitive scores in drug-naïve, first episode SCZ patients compared to HCs [68].Here we found that OLA treatment decreased cecal acetate levels in female SCZ-FMT mice only, and decreased cecal propionic acid levels in both HC-FMT and SCZ-FMT female mice, but to a much greater extent in SCZ-FMT female mice. While these results seem in contrast with the literature, they might be associated with the females’ susceptibility towards developing hyperglycemia, and insulin resistance [70, 71], as only SCZ-FMT female mice presented with increased insulin resistance. In fact, in male mice OLA increased cecal acetic acid levels irrespective of microbiota but did not impact insulin resistance or glycemia. Acetate’s role in regulating glucose metabolism may explain this effect [70, 71], which could protect male mice against OLA-induced hyperglycemia and insulin resistance. Additionally, SCZ-FMT increased cecal isovaleric acid levels in both male and female mice. Isovaleric acid is a branched-SCFA that has been associated with diabetes mellitus [71], and most importantly, higher fecal isovaleric acid levels have been associated with subclinical inflammation, poor cognitive performance [25], and aggression [72] in SCZ patients.

These alterations in the metabolic parameters and fecal SCFAs levels are likely to be consequences of altered GMB composition. We found that GMB composition of the human donors and the recipient mice showed close clustering, suggesting a successful human-to-mouse microbiota transplantation. SCZ microbiota i.e., SCZ-FMT, showed different microbial composition in both male and female mice compared to HC-FMT, suggesting disease (SCZ)-specific changes in the GMB composition. This is in line with previous preclinical [37, 38] and clinical findings reporting distinct microbial composition between SCZ patients and HCs [26, 27, 38, 73, 74]. In line with previous preclinical findings, OLA treatment impacted the microbial composition of SCZ-FMT mice [32, 34, 75, 76]. In female SCZ-FMT mice we found a lower microbial diversity, while OLA treatment increased it. Previous studies show similar results but in male mice [37, 38, 43]. Previous studies have shown increased abundance of the phylum Firmicutes and a decrease in Bacteroidetes abundance in SCZ patients [17] and rodent model [77]. Additionally, APs tend to cause an obesogenic shift and increase the Firmicutes/Bacteroidetes (F/B) ratio in SCZ patients which has been associated with metabolic weight gain [17, 33]. Here we found that SCZ-FMT had an increased F/B ratio, but only in female mice. These results might be linked to the sex-specific glucose dysregulation seen exclusively in female mice and might be related to the differential regulation of acetate and propionate observed only in female SCZ-FMT mice. While OLA increased the F/B ratio in HC-FMT male mice, this change in the microbiome did not appear to influence weight gain, glucose metabolism, or SCFAs levels.

In agreement with previous clinical findings [26, 27, 38, 73, 74], and preclinical studies [37, 38, 43] we found several bacterial genera differentially abundant in SCZ-FMT mice, when compared to HC-FMTT, such as *Lachnnospiraceae* and *Oscillospiraceae*.

APs, mainly OLA and risperidone, alter microbial composition in rodents [32, 34, 75, 78, 79] and in SCZ patient [17, 58, 63, 80], which has been greatly associated with AP-induced metabolic alterations [17]. OLA’s antimicrobial effect also directly impacts microbial composition [32]. Here we found that OLA treatment increased the relative abundance of *Anaerotruncus*, NK4A136 group, *Blautia* and *Parasutterella* species, among others, in SCZ-FMT female mice while it increased *Escherichia-Shigella, Faecalitalea*, *Flavonifractor*, and *Anaerofustis* in male SCZ-FMT mice (Fig. 5C, D). Altogether, OLA increased the relative abundance of many Firmicutes members in both male and female SCZ-FMT mice, which has been associated with AP-induced metabolic alterations [32, 34, 75, 79].

To assess whether compositional microbial differences induced by OLA treatment in SCZ-FMT mice corresponded to functional shifts, we analyzed functional modules and pathways relevant to SCZ pathophysiology. Indeed, Female SCZ-FMT mice showed enrichment of modules involved in glutamate-glutamine-GABA cycle (Fig. 6A). These findings are in agreement with previous SCZ-FMT studies in male mice [20, 43]. Female SCZ-FMT mice also presented with an increase in microbial metabolic pathways related to the synthesis and catabolism of fatty acids, gluconeogenesis, and butyrate synthesis and metabolism, and OLA treatment increased microbial metabolic pathways related to butyrate synthesis and metabolism, and synthesis of glutamine from glutamate.

To summarize, SCZ microbiota induced insulin resistance in female mice which associated with reduced microbial diversity, an increased F/B ratio, and a gut microbiome distinct both at the taxonomic level and at the functional level (SCFAs and other inferred microbial metabolic pathways). Chronic olanzapine treatment induced hyperglycemia only in SCZ-FMT female mice. However, the already obesogenic vehicle (control) (HFD) diet might have potentially masked additional metabolic perturbations induced by chronic olanzapine treatment. Further investigations, including a different vehicle diet (control) and a longer treatment, are warranted. The present study lays the groundwork for future investigations into the role of SCZ-FMT, which will shed light on the GMB’s function in SCZ and the inherent risks of T2D, in addition to guiding the development of animal models and pharmacotherapeutic interventions.

## Conflicts of interests

### Authors contributions

**Raghunath Singh:** Conceptualization; Data curation; Formal analysis; Writing – original draft

**Sandra Pereira:** Writing – review & editing

**Kristoffer Panganiban:** Writing – review & editing

**Laurie Hamel:** Writing – review & editing

**Thomas D. Prevot:** Writing – review & editing

**Daniel J. Mueller:** Writing – review & editing

**Gary Remington:** Writing – review & editing

**Sri Mahavir Agarwal:** Writing – review & editing

**David C. Wright:** Writing – review & editing

**Elena F. Verdu:** Writing – review & editing

**Premysl Bercik:** Writing – review & editing

**Giada De Palma:** Writing – review & editing; Formal analysis. Data generation. Supervision; Validation.

**Margaret K. Hahn:** Conceptualization; Writing – review & editing, Supervision; Validation; Funding acquisition.

### Funding acknowledgments

RS was supported by CIHR postdoctoral fellowship (2023-2026), BBDC’s D. H. Gales Family Charitable Foundation Postdoctoral Fellowship (2022-2023), Bebensee Schizophrenia research postdoctoral fellowship (2020-2022).

This project was funded by BBDC Pilot and feasibility grant 2021 awarded to MKH. MKH is also supported by the CAMH and University of Toronto Meighen Family Research Chair.

GR has received research support from the CIHR, University of Toronto, and HLS Therapeutics Inc.

EFV holds a Canada Research Chair (CIHR) Tier 1 in Microbial Therapeutics and Nutrition in Gastroenterology and is supported by CIHR;

PB holds a Richard Hunt-AstraZeneca Chair in Gastroenterology and is supported by CIHR; GDP is supported by CIHR and Farncombe Innovation Fund.

---

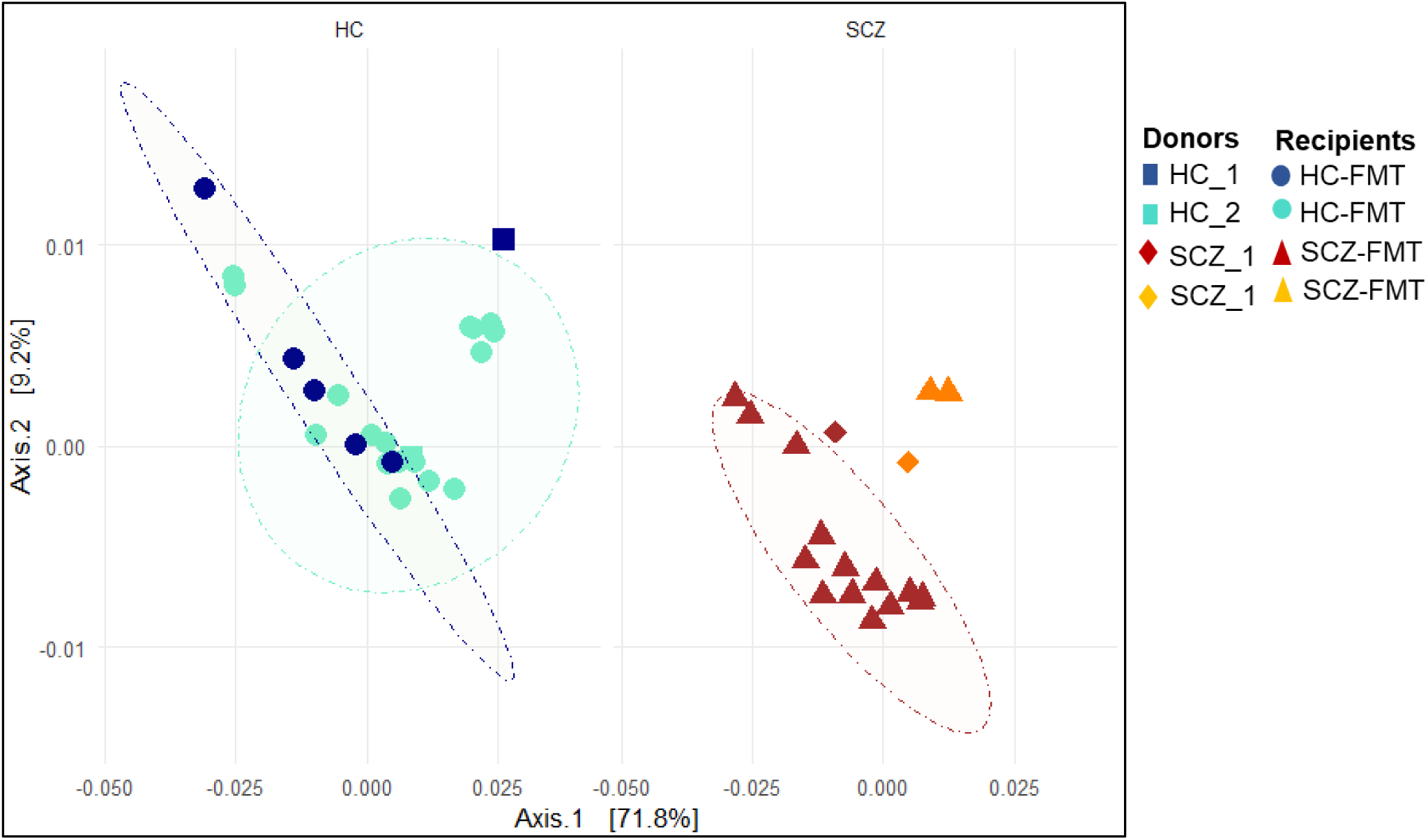
Supplementary figure 1.

## References

1. Marder SR, Cannon TD. Schizophrenia. N Engl J Med. 2019;381:1753–1761.

2. McGrath J, Saha S, Chant D, Welham J. Schizophrenia: A concise overview of incidence, prevalence, and mortality. Epidemiol Rev. 2008.

3. Olfson M, Gerhard T, Huang C, Crystal S, Stroup TS. Premature mortality among adults with schizophrenia in the United States. JAMA Psychiatry. 2015. 2015. 10.1001/jamapsychiatry.2015.1737.

4. Papanastasiou E. The prevalence and mechanisms of metabolic syndrome in schizophrenia: a review. Ther Adv Psychopharmacol. 2013;3:33–51.

5. Perry BI, McIntosh G, Weich S, Singh S, Rees K. The association between first-episode psychosis and abnormal glycaemic control: systematic review and meta-analysis. The Lancet Psychiatry. 2016;3:1049–1058.

6. Perry BI, Stochl J, Upthegrove R, Zammit S, Wareham N, Langenberg C, et al. Longitudinal Trends in Childhood Insulin Levels and Body Mass Index and Associations With Risks of Psychosis and Depression in Young Adults. JAMA Psychiatry. 2021;78:416.

7. Singh R, Bansal Y, Medhi B, Kuhad A. Antipsychotics-induced metabolic alterations: Recounting the mechanistic insights, therapeutic targets and pharmacological alternatives. Eur J Pharmacol. 2019;844:231–240.

8. Shamshoum H, Medak KD, Townsend LK, Ashworth KE, Bush ND, Hahn MK, et al. AMPK β1 activation suppresses antipsychotic-induced hyperglycemia in mice. FASEB J. 2019;33:14010–14021.

9. Castellani LN, Peppler WT, Sutton CD, Whitfield J, Charron MJ, Wright DC. Glucagon receptor knockout mice are protected against acute olanzapine-induced hyperglycemia. Psychoneuroendocrinology. 2017;82:38–45.

10. Medak KD, Townsend LK, Hahn MK, Wright DC. Female mice are protected against acute olanzapine-induced hyperglycemia. Psychoneuroendocrinology. 2019;110:104413.

11. Castellani LN, Wilkin J, Abela A, Benarroch L, Ahasan Z, Teo C, et al. Effects of acute olanzapine exposure on central insulin-mediated regulation of whole body fuel selection and feeding. Psychoneuroendocrinology. 2018;98:127–130.

12. Kowalchuk C, Castellani LN, Chintoh A, Remington G, Giacca A, Hahn MK. Antipsychotics and glucose metabolism: How brain and body collide. Am J Physiol – Endocrinol Metab. 2019;316:E1–E15.

13. Castellani LN, Pereira S, Kowalchuk C, Asgariroozbehani R, Singh R, Wu S, et al. Antipsychotics impair regulation of glucose metabolism by central glucose. Mol Psychiatry. 2022:1–13.

14. Pereira S, Castellani LN, Kowalchuk C, Alganem K, Zhang X, Ryan WG, et al. Olanzapine’s effects on hypothalamic transcriptomics and kinase activity. Psychoneuroendocrinology. 2024;163:106987.

15. Kowalchuk C, Kanagasundaram P, Belsham DD, Hahn MK. Antipsychotics differentially regulate insulin, energy sensing, and inflammation pathways in hypothalamic rat neurons. Psychoneuroendocrinology. 2019;104:42–48.

16. Kowalchuk C, Kanagasundaram P, McIntyre WB, Belsham DD, Hahn MK. Direct effects of antipsychotic drugs on insulin, energy sensing and inflammatory pathways in hypothalamic mouse neurons. Psychoneuroendocrinology. 2019;109:104400.

17. Singh R, Stogios N, Smith E, Lee J, Maksyutynsk K, Au E, et al. Gut microbiome in schizophrenia and antipsychotic-induced metabolic alterations: a scoping review. Ther Adv Psychopharmacol. 2022;12:204512532210965.

18. Nikolova VL, Hall MRB, Hall LJ, Cleare AJ, Stone JM, Young AH. Perturbations in gut microbiota composition in psychiatric disorders: a review and meta-analysis. JAMA Psychiatry. 2021;78:1343–1354.

19. Wang Z, Yuan X, Zhu Z, Pang L, Ding S, Li X, et al. Multiomics analyses reveal microbiome–gut–brain crosstalk centered on aberrant gamma-aminobutyric acid and tryptophan metabolism in drug-naïve patients with first-episode schizophrenia. Schizophr Bull. 2024;50:187–198.

20. Zhu F, Ju Y, Wang W, Wang Q, Guo R, Ma Q, et al. Metagenome-wide association of gut microbiome features for schizophrenia. Nat Commun. 2020;11:1–10.

21. Schwarz E, Maukonen J, Hyytiäinen T, Kieseppä T, Orešič M, Sabunciyan S, et al. Analysis of microbiota in first episode psychosis identifies preliminary associations with symptom severity and treatment response. Schizophr Res. 2018;192:398–403.

22. Vasileva SS, Yang Y, Baker A, Siskind D, Gratten J, Eyles D. Associations of the Gut Microbiome With Treatment Resistance in Schizophrenia. JAMA Psychiatry. 2024. 31 January 2024. 10.1001/jamapsychiatry.2023.5371.

23. He Y, Kosciolek T, Tang J, Zhou Y, Li Z, Ma X, et al. Gut microbiome and magnetic resonance spectroscopy study of subjects at ultra-high risk for psychosis may support the membrane hypothesis. Eur Psychiatry. 2018;53:37–45.

24. Zhu C, Zheng M, Ali U, Xia Q, Wang Z, Chenlong, et al. Association Between Abundance of Haemophilus in the Gut Microbiota and Negative Symptoms of Schizophrenia. Front Psychiatry. 2021;12:1230.

25. Kowalski K, Szponar B, Bochen P, Żebrowska-Różańska P, Łaczmański Ł, Samochowiec J, et al. Altered levels of fecal short-chain fatty acids are associated with subclinical inflammation and worse cognitive performance in patients with schizophrenia. J Psychiatr Res. 2023;165:298–304.

26. Thirion F, Speyer H, Hansen TH, Nielsen T, Fan Y, Le Chatelier E, et al. Alteration of gut microbiome in patients with schizophrenia indicates links between bacterial tyrosine biosynthesis and cognitive dysfunction. Biol Psychiatry Glob Open Sci. 2023;3:283–291.

27. Stiernborg M, Prast-Nielsen S, Melas PA, Skott M, Millischer V, Boulund F, et al. Differences in the gut microbiome of young adults with schizophrenia spectrum disorder: using machine learning to distinguish cases from controls. Brain Behav Immun. 2024;117:298–309.

28. Ling Z, Lan Z, Cheng Y, Liu X, Li Z, Yu Y, et al. Altered gut microbiota and systemic immunity in Chinese patients with schizophrenia comorbid with metabolic syndrome. J Transl Med. 2024;22:729.

29. Xu Y, Shao M, Fang X, Tang W, Zhou C, Hu X, et al. Antipsychotic-induced gastrointestinal hypomotility and the alteration in gut microbiota in patients with schizophrenia. Brain Behav Immun. 2022;99:119–129.

30. Ma X, Asif H, Dai L, He Y, Zheng W, Wang D, et al. Alteration of the gut microbiome in first-episode drug-naïve and chronic medicated schizophrenia correlate with regional brain volumes. J Psychiatr Res. 2020;123:136–144.

31. Dinan TG, Cryan JF. Schizophrenia and the microbiome: Time to focus on the impact of antipsychotic treatment on the gut microbiota. World J Biol Psychiatry. 2018;19:568–570.

32. Morgan AP, Crowley JJ, Nonneman RJ, Quackenbush CR, Miller CN, Ryan AK, et al. The antipsychotic olanzapine interacts with the gut microbiome to cause weight gain in mouse. PLoS One. 2014;9:e115225.

33. Bahr SM, Weidemann BJ, Castro AN, Walsh JW, DeLeon O, Burnett CML, et al. Risperidone-induced weight gain is mediated through shifts in the gut microbiome and suppression of energy expenditure. EBioMedicine. 2015;2:1725–1734.

34. Davey KJ, Cotter PD, O’Sullivan O, Crispie F, Dinan TG, Cryan JF, et al. Antipsychotics and the gut microbiome: olanzapine-induced metabolic dysfunction is attenuated by antibiotic administration in the rat. Transl Psychiatry. 2013;3:1–7.

35. Kao AC-C, Spitzer S, Anthony DC, Lennox B, Burnet PWJ. Prebiotic attenuation of olanzapine-induced weight gain in rats: analysis of central and peripheral biomarkers and gut microbiota. Transl Psychiatry. 2018;8:1–12.

36. Singh R, Panganiban K, Au E, Ravikumar R, Pereira S, Prevot TD, et al. Human-fecal microbiota transplantation in relation to gut microbiome signatures in animal models for schizophrenia: A scoping review. Asian J Psychiatr. 2024:104285.

37. Zhu F, Guo R, Wang W, Ju Y, Wang Q, Ma Q, et al. Transplantation of microbiota from drug-free patients with schizophrenia causes schizophrenia-like abnormal behaviors and dysregulated kynurenine metabolism in mice. Mol Psychiatry. 2020;25:2905–2918.

38. Wei N, Ju M, Su X, Zhang Y, Huang Y, Rao X, et al. Transplantation of gut microbiota derived from patients with schizophrenia induces schizophrenia-like behaviors and dysregulated brain transcript response in mice. Schizophrenia. 2024;10:44.

39. De Palma G, Lynch MDJ, Lu J, Dang VT, Deng Y, Jury J, et al. Transplantation of fecal microbiota from patients with irritable bowel syndrome alters gut function and behavior in recipient mice. Sci Transl Med. 2017;9.

40. De Palma G, Shimbori C, Reed DE, Yu Y, Rabbia V, Lu J, et al. Histamine production by the gut microbiota induces visceral hyperalgesia through histamine 4 receptor signaling in mice. Sci Transl Med. 2022;14:eabj1895.

41. Lord CC, Wyler SC, Wan R, Castorena CM, Ahmed N, Mathew D, et al. The atypical antipsychotic olanzapine causes weight gain by targeting serotonin receptor 2C. J Clin Invest. 2017;127:3402–3406.

42. Zhao S, Lin Q, Xiong W, Li L, Straub L, Zhang D, et al. Hyperleptinemia contributes to antipsychotic drug–associated obesity and metabolic disorders. Sci Transl Med. 2023;15:eade8460.

43. Zheng P, Zeng B, Liu M, Chen J, Pan J, Han Y, et al. The gut microbiome from patients with schizophrenia modulates the glutamate-glutamine-GABA cycle and schizophrenia-relevant behaviors in mice. Sci Adv. 2019;5:eaau8317.

44. Whelan FJ, Surette MG. A comprehensive evaluation of the sl1p pipeline for 16S rRNA gene sequencing analysis. Microbiome. 2017;5:1–13.

45. Bartram AK, Lynch MDJ, Stearns JC, Moreno-Hagelsieb G, Neufeld JD. Generation of multimillion-sequence 16S rRNA gene libraries from complex microbial communities by assembling paired-end Illumina reads. Appl Environ Microbiol. 2011;77:3846–3852.

46. Callahan BJ, McMurdie PJ, Rosen MJ, Han AW, Johnson AJA, Holmes SP. DADA2: High-resolution sample inference from Illumina amplicon data. Nat Methods. 2016. 2016. 10.1038/nmeth.3869.

47. McMurdie PJ, Holmes S. Phyloseq: An R Package for Reproducible Interactive Analysis and Graphics of Microbiome Census Data. PLoS One. 2013. 2013. 10.1371/journal.pone.0061217.

48. Wang Q, Garrity GM, Tiedje JM, Cole JR. Naïve Bayesian classifier for rapid assignment of rRNA sequences into the new bacterial taxonomy. Appl Environ Microbiol. 2007. 2007. 10.1128/AEM.00062-07.

49. Quast C, Pruesse E, Yilmaz P, Gerken J, Schweer T, Yarza P, et al. The SILVA ribosomal RNA gene database project: improved data processing and web-based tools. Nucleic Acids Res. 2012;41:D590–D596.

50. Yilmaz P, Parfrey LW, Yarza P, Gerken J, Pruesse E, Quast C, et al. The SILVA and “All-species Living Tree Project (LTP)” taxonomic frameworks. Nucleic Acids Res. 2014;42:D643–D648.

51. Bolyen E, Rideout JR, Dillon MR, Bokulich NA, Abnet CC, Al-Ghalith GA, et al. Reproducible, interactive, scalable and extensible microbiome data science using QIIME 2. Nat Biotechnol. 2019;37:852–857.

52. Mallick H, Rahnavard A, McIver LJ, Ma S, Zhang Y, Nguyen LH, et al. Multivariable association discovery in population-scale meta-omics studies. PLOS Comput Biol. 2021;17:e1009442.

53. Douglas GM, Maffei VJ, Zaneveld JR, Yurgel SN, Brown JR, Taylor CM, et al. PICRUSt2 for prediction of metagenome functions. Nat Biotechnol. 2020;38:685–688.

54. Parks DH, Tyson GW, Hugenholtz P, Beiko RG. STAMP: statistical analysis of taxonomic and functional profiles. Bioinformatics. 2014;30:3123–3124.

55. Albaugh VL, Vary TC, Ilkayeva O, Wenner BR, Maresca KP, Joyal JL, et al. Atypical antipsychotics rapidly and inappropriately switch peripheral fuel utilization to lipids, impairing metabolic flexibility in rodents. Schizophr Bull. 2012. 2012. 10.1093/schbul/sbq053.

56. Hahn MK, Chintoh A, Remington G, Teo C, Mann S, Arenovich T, et al. Effects of intracerebroventricular (ICV) olanzapine on insulin sensitivity and secretion in vivo: An animal model. Eur Neuropsychopharmacol. 2014;24:448–458.

57. Singh R, Bansal Y, Sodhi RK, Saroj P, Medhi B, Kuhad A. Modeling of antipsychotic-induced metabolic alterations in mice: An experimental approach precluding psychosis as a predisposing factor. Toxicol Appl Pharmacol. 2019;378:1–13.

58. Bahr SM, Tyler BC, Wooldridge N, Butcher BD, Burns TL, Teesch LM, et al. Use of the second-generation antipsychotic, risperidone, and secondary weight gain are associated with an altered gut microbiota in children. Transl Psychiatry. 2015;5:e652.

59. Zapata RC, Osborn O. Susceptibility of male wild type mouse strains to antipsychotic-induced weight gain. Physiol Behav. 2020;220:112859.

60. Soto PL, Young ME, Nguyen S, Federoff M, Goodson M, Morrison CD, et al. Early adolescent second-generation antipsychotic exposure produces long-term, post-treatment increases in body weight and metabolism-associated gene expression. Pharmacol Biochem Behav. 2024:173951.

61. Yuan X, Zhang Y, Pang L, Zhang X, Kang Y, Hei G, et al. Insulin Resistance Links Dysbiosis of Gut Microbiota with Cognitive Impairment in First-Episode Drug-Naïve Schizophrenia. Psychoneuroendocrinology. 2024:107255.

62. Misiak B, Pawlak E, Rembacz K, Kotas M, Żebrowska-Różańska P, Kujawa D, et al. Associations of gut microbiota alterations with clinical, metabolic, and immune-inflammatory characteristics of chronic schizophrenia. J Psychiatr Res. 2024;171:152– 160.

63. Li X, Yuan X, Pang L, Miao Y, Wang S, Zhang X, et al. Gut Microbiota Markers for Antipsychotics Induced Metabolic Disturbance in Drug Naïve Patients with First Episode Schizophrenia–A 24 Weeks Follow-up Study. MedRxiv. 2021:2012–2020.

64. Yuan X, Zhang P, Wang Y, Liu Y, Li X, Kumar BU, et al. Changes in metabolism and microbiota after 24-week risperidone treatment in drug naïve, normal weight patients with first episode schizophrenia. Schizophr Res. 2018;201:299–306.

65. Dalile B, Van Oudenhove L, Vervliet B, Verbeke K. The role of short-chain fatty acids in microbiota–gut–brain communication. Nat Rev Gastroenterol Hepatol. 2019;16:461–478.

66. Peng H, Ouyang L, Li D, Li Z, Yuan L, Fan L, et al. Short-chain fatty acids in patients with schizophrenia and ultra-high risk population. Front Psychiatry. 2022;13:977538.

67. González Hernández MA, Canfora EE, Jocken JWE, Blaak EE. The short-chain fatty acid acetate in body weight control and insulin sensitivity. Nutrients. 2019;11:1943.

68. Li X, Yuan X, Pang L, Zhang S, Li Y, Huang X, et al. The effect of serum lipids and short-chain fatty acids on cognitive functioning in drug-naïve, first episode schizophrenia patients. Psychiatry Res. 2022;313:114582.

69. Tirosh A, Calay ES, Tuncman G, Claiborn KC, Inouye KE, Eguchi K, et al. The short-chain fatty acid propionate increases glucagon and FABP4 production, impairing insulin action in mice and humans. Sci Transl Med. 2019;11:eaav0120.

70. He Q, Ji L, Wang Y, Zhang Y, Wang H, Wang J, et al. Acetate enables metabolic fitness and cognitive performance during sleep disruption. Cell Metab. 2024;36:1998–2014.

71. Wijdeveld M, Schrantee A, Hagemeijer A, Nederveen AJ, Scheithauer TPM, Levels JHM, et al. Intestinal acetate and butyrate availability is associated with glucose metabolism in healthy individuals. Iscience. 2023;26.

72. Deng H, He L, Wang C, Zhang T, Guo H, Zhang H, et al. Altered gut microbiota and its metabolites correlate with plasma cytokines in schizophrenia inpatients with aggression. BMC Psychiatry. 2022;22:629.

73. Yuan X, Wang Y, Li X, Jiang J, Kang Y, Pang L, et al. Gut microbial biomarkers for the treatment response in first-episode, drug-naïve schizophrenia: a 24-week follow-up study. Transl Psychiatry. 2021;11:422.

74. Pan R, Zhang X, Gao J, Yi W, Wei Q, Su H. Analysis of the diversity of intestinal microbiome and its potential value as a biomarker in patients with schizophrenia: a cohort study. Psychiatry Res. 2020;291:113260.

75. Davey KJ, O’Mahony SM, Schellekens H, O’Sullivan O, Bienenstock J, Cotter PD, et al. Gender-dependent consequences of chronic olanzapine in the rat: effects on body weight, inflammatory, metabolic and microbiota parameters. Psychopharmacology (Berl). 2012;221:155–169.

76. Kamath S, Hunter A, Collins K, Wignall A, Joyce P. The atypical antipsychotics lurasidone and olanzapine exert contrasting effects on the gut microbiome and metabolic function of rats. Br J Pharmacol. 2024;181:4531–4545.

77. Juckel G, Manitz M-P, Freund N, Gatermann S. Impact of Poly I: C induced maternal immune activation on offspring’s gut microbiome diversity–implications for schizophrenia. Prog Neuro-Psychopharmacology Biol Psychiatry. 2021;110:110306.

78. Bahr SM, Weidemann BJ, Castro AN, Walsh JW, DeLeon O, Burnett CML, et al. Risperidone-induced weight gain is mediated through shifts in the gut microbiome and suppression of energy expenditure. EBioMedicine. 2015;2:1725–1734.

79. Qian L, He X, Liu Y, Gao F, Lu W, Fan Y, et al. Longitudinal Gut Microbiota Dysbiosis Underlies Olanzapine-Induced Weight Gain. Microbiol Spectr. 2023;11:e00058–23.

80. Flowers SA, Baxter NT, Ward KM, Kraal AZ, McInnis MG, Schmidt TM, et al. Effects of Atypical Antipsychotic Treatment and Resistant Starch Supplementation on Gut Microbiome Composition in a Cohort of Patients with Bipolar Disorder or Schizophrenia. Pharmacotherapy. 2019;39:161–170.

